# Unsupervised Contrastive Peak Caller for ATAC-seq

**DOI:** 10.1101/2023.01.07.523108

**Authors:** Ha T.H. Vu, Yudi Zhang, Geetu Tuteja, Karin Dorman

## Abstract

The assay for transposase-accessible chromatin with sequencing (ATAC-seq) is a common assay to identify chromatin accessible regions by using a Tn5 transposase that can access, cut, and ligate adapters to DNA fragments for subsequent amplification and sequencing. These sequenced regions are quantified and tested for enrichment in a process referred to as “peak calling”. Most unsupervised peak calling methods are based on simple statistical models and suffer from elevated false positive rates. Newly developed supervised deep learning methods can be successful, but they rely on high quality labeled data for training, which can be difficult to obtain. Moreover, though biological replicates are recognized to be important, there are no established approaches for using replicates in the deep learning tools, and the approaches available for traditional methods either cannot be applied to ATAC-seq, where control samples may be unavailable, or are post-hoc and do not capitalize on potentially complex, but reproducible signal in the read enrichment data. Here, we propose a novel peak caller that uses unsupervised contrastive learning to extract shared signals from multiple replicates. Raw coverage data are encoded to obtain low-dimensional embeddings and optimized to minimize a contrastive loss over biological replicates. These embeddings are passed to another contrastive loss for learning and predicting peaks and decoded to denoised data under an autoencoder loss. We compared our Replicative Contrastive Learner (RCL) method with other existing methods on ATAC-seq data, using annotations from ChromHMM genome and transcription factor ChIP-seq as noisy truth. RCL consistently achieved the best performance.

---

The assay for transposase-accessible chromatin with sequencing (ATAC-seq) is widely used when studying chromatin biology (Grandi et al. 2022). ATAC-seq utilizes a hyperactive mutant Tn5 transposase to cleave double stranded DNA and to attach adapters for subsequent sequencing by high throughput technologies (Buenrostro et al. 2015). Since DNA is more easily cleaved where it is unwound and open, sequenced DNA fragments tend to arise from regions of open chromatin. A standard analysis for ATAC-seq starts with aligning the sequencing reads to a reference genome using BWA (H. Li 2013), Bowtie2 (Langmead and Salzberg 2012), or other short read aligner (Musich et al. 2021). Then peak calling methods will identify the open regions (peaks) in the genome where aligned reads are enriched. Downstream analyses include motif detection, differential binding analysis or footprint identification (Buenrostro et al. 2013; Grandi et al. 2022), all of which require accurate peak calls. Unfortunately, peaks of false enrichment may be called due to mapping errors or experimental noise (Park 2009). Such errors can be reduced by masking repetitive regions and using control samples (Zhang et al. 2008), but input controls for ATAC-seq are typically not used due to high sequencing costs (F. Yan et al. 2020).

ATAC-seq peaks are often called with the most popular general-purpose peak caller, MACS (Zhang et al. 2008), and there is an ATAC-seq-specific method called HMMRATAC (Tarbell and Liu 2019). MACS slides a fixed-width window across the genome to find candidate peaks. The number of reads aligned to the genome in the current window is modeled as a Poisson random variable, with a dynamic mean to capture local variation in background coverage rates. MACS calculates the *p*-value for each candidate peak as the probability of obtaining coverage at or above the observed coverage given the current background rate. HMMRATAC (Tarbell and Liu 2019) employs a hidden Markov model (HMM) with four-dimensional emissions of varying fragment sizes, nucleosome-free (NF), one nucleosome (1N), two nucleosome (2N) and three nucleosome (3N) fragments, from three possible hidden states: a “center” state (open chromatin), with high emissions in all four dimensions, a nucleosome state, with low NF fragment emission, and a background state, with low emissions in all dimensions. Once the HMM has been estimated, the Viterbi algorithm is used to classify every 10 base pair (bp) window in the genome into one of the three states.

Traditional modeling methods tend to predict many false positive peaks in ChIP-seq applications (Hocking et al. 2017), and some investigations have shown humans to be superior “peak callers” (Rye et al. 2011; Hocking et al. 2017). Inspired by such human performance and recent successes in artificial intelligence, two new peak callers, CNN-Peaks (Oh et al. 2020) and LanceOtron (Hentges et al. 2021), take a deep learning approach. CNN-Peaks (Oh et al. 2020) uses supervised convolutional neural networks (CNN) to call ChIP-seq peaks. In addition to the read count information obtained from BAM files, it uses genome annotation information, such as protein-coding transcripts, to improve estimation of peak locations. In their CNN architecture, filters of various sizes are used to extract diverse features and a weighted cross-entropy loss is adopted to account for the imbalanced labels. LanceOtron (Hentges et al. 2021) is another supervised CNN based deep learning method that can be used on ATAC-seq, ChIP-seq, and DNase-seq data. It feeds the output of a logistic regression, fit to 11 enrichment scores predicting labeled peaks, the output of a CNN, fit to fragment coverage in 2000 bp windows predicting labeled peaks, and the 11 enrichment scores to a multilayer perceptron to produce the overall peak score. Many of the false-positive peaks generated by other peak callers are filtered out by these supervised deep learners, increasing precision by about 18% (Hentges et al. 2021). Unfortunately, these supervised methods require labeled data for model training, which are often hard or costly to obtain.

None of these methods consider biological replicates, and in fact most peak calling methods assess biological replicates separately (Goren et al. 2018). HMMRATAC and some users of MACS recommend combining multiple replicates to increase signal, but joint analysis of multiple biological replicates could improve the power to distinguish actual transcription factor binding events (Newell et al. 2021), since some weak or highly variable peak signals may only become evident across multiple replicates (Zhang et al. 2014). One common approach for assessing reproducibility from replicates uses the Irreproducible Discovery Rate (IDR), which identifies reproducible peaks by measuring the consistency in peak ranks between replicates (Q. Li et al. 2011). ChIP-R (Newell et al. 2021), which shows improvement over IDR and can handle more than two replicates, uses the rank product to evaluate the reproducibility across any number of ChIP-seq or ATAC-seq replicates.

We propose a novel unsupervised learning method that uses contrastive learning (Le-Khac et al. 2020) across replicates to separate genomic regions into peaks and non-peaks. This method overcomes excess noise and lack of labels to make better inferences than existing methods. The entire pipeline is released under the GNU Public License to the community as a package named RCL, for Replicative Contrastive Learner (GitHub https://github.com/Tuteja-Lab/UnsupervisedPeakCaller).

## 1 Results

### 1.1 The RCL algorithm

In this study, we developed a peak calling tool (RCL), which contrasts biological replicates to identify the shared signals of ATAC-seq peaks (Figure 1). Peak calling is a difficult task, where the genomic extent and significance of enrichment, together the

**Figure 1:**
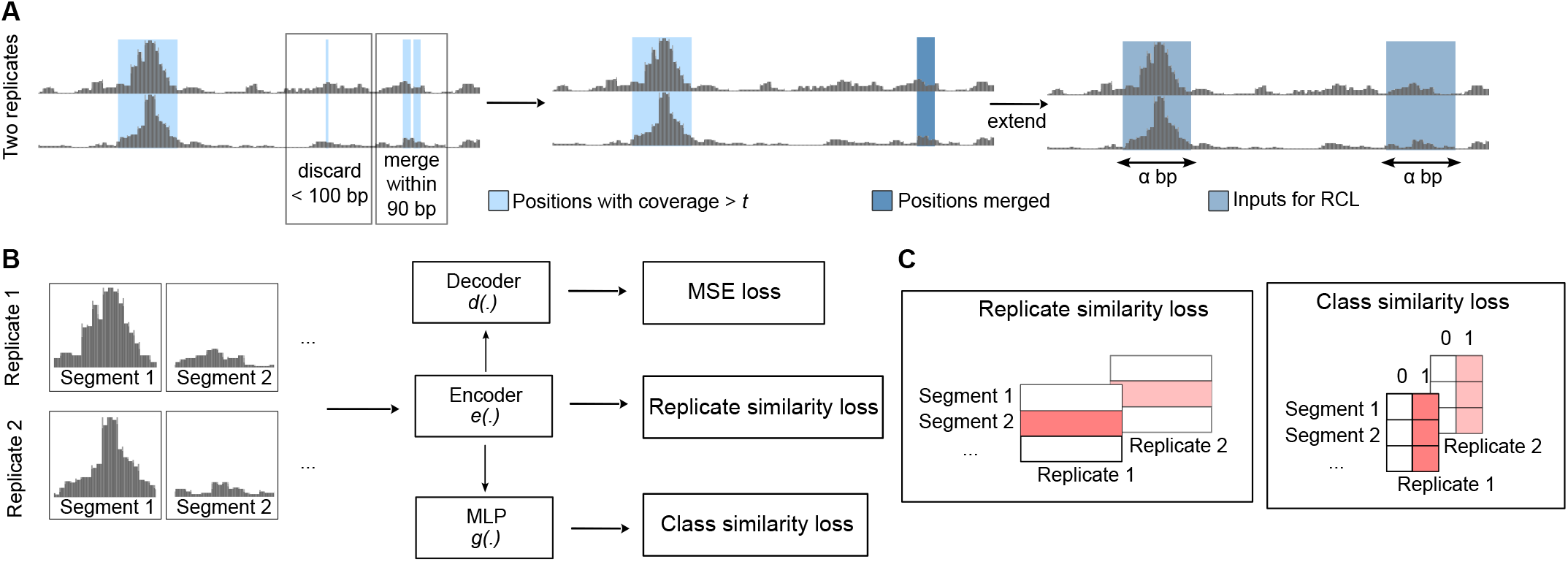
RCL model. Segment identifies the same genomic region for all replicates. (**A**) The raw input is processed to extract *α*-length input segments. (**B**) The *α*-length input segments are fed to encoder *e*(·) to compute the cross-replicate contrastive loss. Then the embedding is fed to a multilayer perceptron (MLP), specifically a fully connected neural network, for class similarity loss and a decoder for the autoencoder (MSE) loss. The encoder/decoder has five ResNET blocks. (**C**) Shaded red boxes represent the elements contrasted in the respective losses.

*peak*, must be inferred. Our proposed method separates these tasks, first liberally identifying candidate regions of possible enrichment, and then learning how to score and classify data extracted from the regions. The learner makes no attempt to learn peak boundaries, so its predictions are passed back to the original candidate regions, which become peak predictions if sufficiently high scoring.

#### 1.1.1 Prediction region selection

The individual BAM files of *R* replicates and a merged BAM file are required to identify candidate peak regions. Additionally, two user settable parameters, coverage threshold (*t*, default: “median”, see Step 1 below) and input segment length (*α*, default: 1,000), affect the number and length of the candidate regions. Given these inputs, candidate peak regions are identified as follows:

**Step 1:** Retain genome positions with coverage *> t* in all *R* individual BAM files. Threshold *t* defaults to a chromosome-specific value obtained from the input data. Specifically, the read coverage on each chromosome is calculated using BEDTools `genomecov` (Quinlan and Hall 2010) for every replicate (`bedtools genomecov –ibam bamFiles -pc -bga`), then median coverage across all nonzero positions per chromosome is obtained. The minimum median observed across replicates for a chromosome is used as the threshold for that chromosome. Alternatively, *t* can be set as a single integer value to be used for every chromosome.

**Step 2:** Contiguous retained sites are aggregated into regions. Then, regions within 90 base pairs (bp) are merged, since DNA linkers are known to be 8–90 bp (Singh and Mueller-Planitz 2021). All regions longer than 100 bp are retained for Step 3. Define this set of regions as 𝒜.

**Step 3.1:** If a region in set 𝒜 is shorter than *α*, an *α* bp long genomic segment is obtained by extending 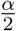 bp up- and down-stream of its midpoint.

**Step 3.2:** For regions in set 𝒜 longer than *α* bp, we first get positions with coverage summed across replicates ≥ 0.95 quantile of the region (obtained from the merged BAM file). Positions within *α* bp are merged, then we extended 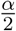 bp up- and down-stream of each merged region’s midpoint.

Hereafter, “segment” refers to these selected *α* bp genomic fragments. Any segment overlapping with a blacklist region (Amemiya et al. 2019) by at least 1 bp is removed. In the end, per-base coverage vectors for these length *α* bp segments from *R* replicates are the inputs to RCL.

#### 1.1.2 Unsupervised learner

Given the candidate peak regions, we use a neural network to assign a score to each segment. These segment scores are combined into a single score for the candidate region (see RCL in §3.4. Method comparison for details), and sufficiently high-scoring regions are called peaks. As illustrated in Fig. 1, our method consists of three jointly learned components along with their respective losses, so the total minimized loss is

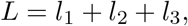

shown without optional weights that can be tuned by standard cross-validation methods. The three components are a cross-replicate contrastive learner (Le-Khac et al. 2020), a segment class (peak/non-peak) learner (Zhong et al. 2020), and an autoencoder (Kramer 1991). The input to the contrastive learner and segment class learner is the output of the encoder network that maps the *α* bp coverage data to a lower-dimensional representation space.

##### Encoder

With *R* replicates of observed coverage in *S* segments, the input data are per-base coverage vectors ***m***_*ri*_, *r* ∈ {1, …, *R*}, *i* ∈ {1, …, *S*}. We use ResNET (He et al. 2016) as the backbone of our encoder network. A ResNET module is composed of three basic blocks followed by one residual block. A basic block is composed of a 1D convolutional layer (default: dilation 8 and kernel size 31), followed by a RELU activation function. Our whole encoder *e*(·) is made of five such ResNET modules, producing the lower dimensional (default dimension: 50) representation ***x***_*ri*_ = *e* (***m***_*ri*_).

##### Replicate-wise contrastive learning

We use the latent space representations ***x***_*ri*_ for computing the cross-replicate contrastive loss. We follow Sim-CLR (Le-Khac et al. 2020), where the replicates are augmentations and the same segments across replicates are positive examples, otherwise they are negative examples. The pairwise replicate contrastive loss, *l*_1_,

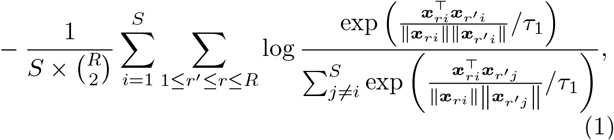

where *τ*_1_ is the temperature hyperparameter (default: 0.5), aims to learn lower-dimensional representations such that positive examples are close and negative examples are distant in the new space.

##### Segment class learning

Assuming the actual peak/non-peak status of genomic segments is shared across replicates and there are underlying characteristics of coverage that define peaks and non-peaks, we expect the low-dimension representation of peak segments to cluster together and separate from the non-peak segments in the new space. Therefore, we also require the representations to match in discrete (classification) space, which we achieve by requiring peak probabilities for each segment to be similar across replicates. The embedded representations ***x***_*ri*_ are reduced to two dimensions via a fully connected neural network (multilayer perceptron, MLP, in the following) with one hidden layer the same dimension as ***x***_*ri*_, followed by the softmax function, together denoted as *g*(·). Letting ***q***_*ri*_ = *g*(***x***_*ri*_) be the peak/non-peak probabilities for segment *i* in replicate *r* and 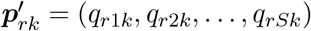, *k* ∈ {1, 2}, vectors of peak/non-peak probabilities across segments for the *r*th replicate, we maximize similarity in peak calls among replicates using loss *l*_2_

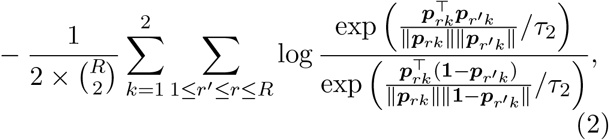

where temperature hyperparameter *τ*_2_ = *τ*_1_ in our experiments. This loss strengthens the shared peak signal across replicates and provides a peak/non-peak prediction for each segment of each replicate.

##### Autoencoder learning

We also want to produce cleaner data in the original genomic space, useful for purposes such as visualization or replicate merging. Therefore, we use decoder *d*(·), with structure symmetric to the encoder *e*(·), to map ***x***_*ri*_ back to predicted data 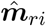 in the genomic space. An autoencoder has good embedded feature representation capability (Baldi 2012), learned by minimizing the squared error loss *l*_3_,

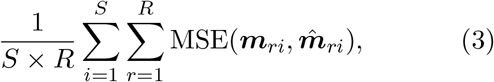

between the original data ***m***_*ri*_ and the reconstructed data 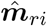.

### 1.2 Performance benchmarking using ChromHMM annotations

We compared the performance of RCL to both unsupervised (MACS, ChIP-R and HMMRATAC) and pre-trained supervised (LanceOtron) peak callers, where we used data from four human cell lines, MCF-7, A549, K652 and GM12878, and one dataset generated from mouse placenta tissues at embryonic day 9.5. The datasets are summarized in Table 1.

**Table 1:** Datasets used to compare methods on genome-wide annotation regions generated by ChromHMM. Src., literature source. Org., organism. *R*, number of biological replicates. Size, mean number of reads across replicates after filtering in the dataset. Map. Rate, median proportion of aligned reads to autosomal and sex chromosomes using Bowtie2. ChromHMM labels, number of positive and negative true regions annotated using ChromHMM. RCL MED (default threshold *t* based on median coverage) and C2 (threshold *t* = 2) indicate two different coverage thresholds used to build training segments. Lower threshold results in larger input datasets and provides more predictions, specifically including predictions for lower coverage, harder-to-predict segments. For the total number of peaks including those outside of annotated regions, see Supplementary Table S1.

| Data | Src. | Org. | $R$ | Size | Map. Rate | ChromHMM Labels | | Number of Called Peaks in ChromHMM Labels | | | | | |
| --- | --- | --- | --- | --- | --- | --- | --- | --- | --- | --- | --- | --- | --- |
|  |  |  |  |  |  | + | - | MACS | ChIP-R | HMMR ATAC | RCL MED | C2 | Lance Otron |
| MCF-7 | Consortium 2012 | Human | 2 | 37M | 96% | 148,531 | 116,729 | 65,440 | 93,860 | 113,362 | 92,333 | 116,367 | 90,050 |
| A549 | Consortium 2012 | Human | 3 | 157M | 98% | 154,274 | 120,726 | 116,654 | 133,121 | 80,991 | 148,725 | 213,263 | 94,284 |
| GM12878 | Buenrostro et al. 2013 | Human | 4 | 31M | 68% | 156,817 | 139,567 | 58,341 | 45,498 | 47,315 | 59,130 | 38,637 | 33,056 |
| K562 | Consortium 2012 | Human | 3 | 100M | 96% | 212,779 | 190,127 | 143,269 | 139,343 | 87,284 | 173,790 | 227,839 | 110,011 |
| Placenta | R. Starks et al. 2019 | Mouse | 3 | 20M | 70% | 88,211 | 18,754 | 14,598 | 15,888 | 6,277 | 11,332 | 5,719 | 4,141 |

The RCL method involves one important tunable parameter – the coverage threshold *t* (option `-t`) used to identify the candidate peak regions and segments for model training. By default, RCL uses a chromosome-specific threshold that depends on the median coverage (see §1.1.1. Prediction region selection). In all datasets, in addition to using this default setting, we also implemented RCL with `-t 2` to explore the impact of this tuning parameter. In datasets with higher library size (MCF-7, K562 and A549), chromosome-specific thresholds generally exceed two (Supplementary Figure S1); default thresholds for lower library size datasets (GM12878 and mouse placenta) for all chromosomes are one. We observed manual lowering of threshold *t* increases the number of candidate regions supplied to the RCL model.

Across all tested datasets of varying library size, RCL achieved the best overall performance (Tables 2a, 2b). Sporadically, HMMRATAC, LanceOtron or ChIP-R achieved higher precision at the cost of much lower recall. As the threshold *t* decreases, RCL predicts more peaks with lower precision and higher recall. Overall, the model trained with lower threshold (RCL-C2 for MCF-7, K562, and A549; RCL-MED for GM12878 and mouse placenta) achieved universally better F1 scores, suggesting that exposure to more low coverage regions can help RCL distinguish true peaks. In these comparisons, MACS and ChIP-R peaks were called with *q*-value 0.05, but HMMRATAC, LanceOtron, and RCL peaks were called without false discovery control. HMMRATAC calls should be filtered by the score (Tarbell and Liu 2019), typically a measure of coverage, and it is similarly advisable to filter RCL calls for higher precision.

**Table 2:**
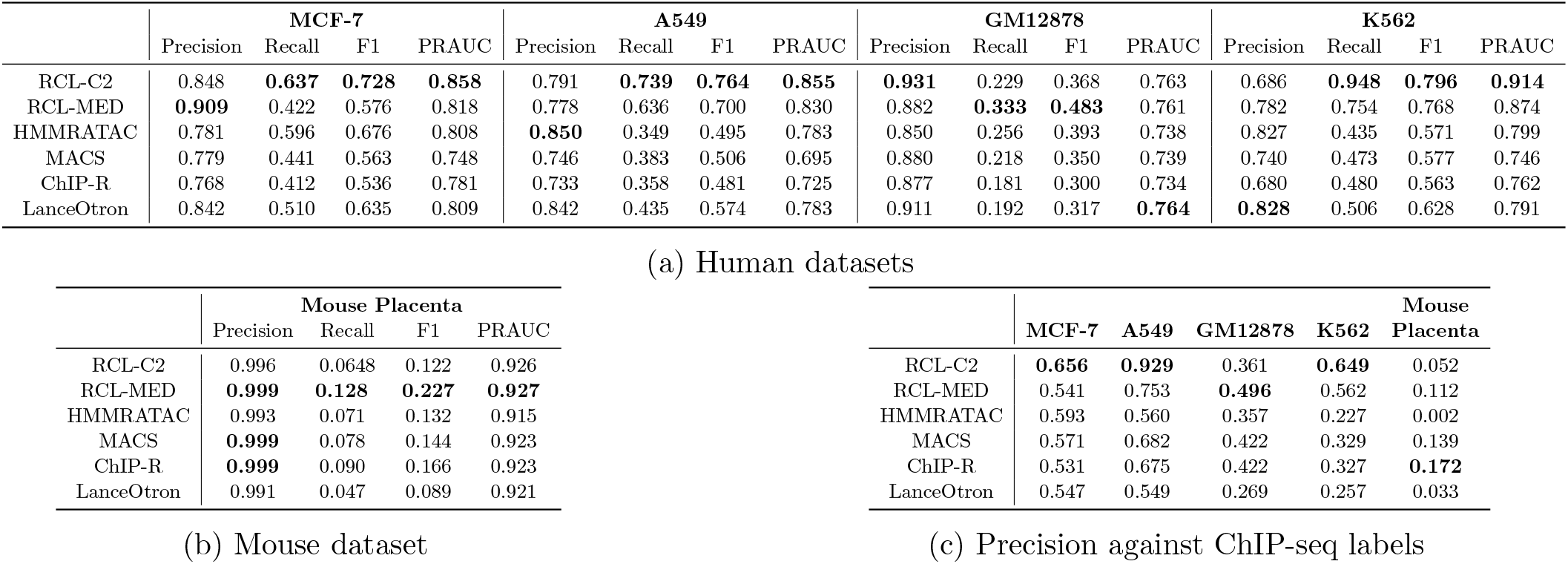
(a) and (b): Precision, recall, F1 scores and PRAUC (area under the PR curve). To compute precision, recall, and F1 scores for MACS and ChIP-R, a *q*-value of 0.05 was used. To compute PRAUC for MACS and ChIP-R, a *q*-value of 0.5 was used and then post-hoc thresholded to obtain a PR curve. All HMMRATAC results were obtained using scores *>* 0. All RCL and LanceOtron results were obtained using average scores across replicates *>* 0.5. (c): Precision using transcription factor (TF) ChIP-seq as labels.

PR curves are useful for comparing methods across all false discovery rates (Figure 2, Supplementary Figure S2). We applied a relaxed *q*-value threshold (0.5) to generate candidate peaks for MACS and ChIP-R, and post-hoc thresholded to plot the curves. Since MACS and ChIP-R are usually run with smaller *q*-values and no post-hoc thresholding, we also plot precision and recall point estimates for typical choices of *q* (methods labeled “multiQ”). The linear portion of each PR curve from the black dot to 100% recall corresponds to the subset of ChromHMM-labeled regions with no score assigned by the method. The RCL PR curve, especially with lower threshold *t*, dominates the curves of other methods. RCL appears to use replicate information in the coverage data better than ChIP-R’s post-hoc comparison of peak calls across replicates, which is generally better than naive aggregation of MACS calls. HMMRATAC achieves intermediate performance (Table 2a, 2b), with higher achievable recall than MACS and ChIP-R of weak peaks, but sometimes lower achievable precision on strong peaks (Figure 2). HMMRATAC also performs poorly on data with lower library size, probably because coverage data are too sparse, when partitioned by fragment length, to estimate this parameter-rich model. Despite the high number of predicted peaks for K562 and A549 data (Table 1), RCL maintained good precision out to much higher recall. In particular, RCL achieved nearly twice as many true predictions while maintaining higher precision than either MACS or LanceOtron. While ChIP-R, and sometimes HMMRATAC, can achieve near equal performance on the strongest peaks, only RCL can maintain high precision on the more difficult peaks. For lower library size GM12878 and mouse data, all methods called a limited number of peaks, with low achievable recall. Nevertheless, RCL still obtained better performance, except at the highest achieved recall, where LanceOtron had higher precision. The slightly lower PRAUC of RCL on these data (Table 2a) may yet be overcome by allowing non-integer threshold values *t* on the average coverage across multiple sites.

**Figure 2:**
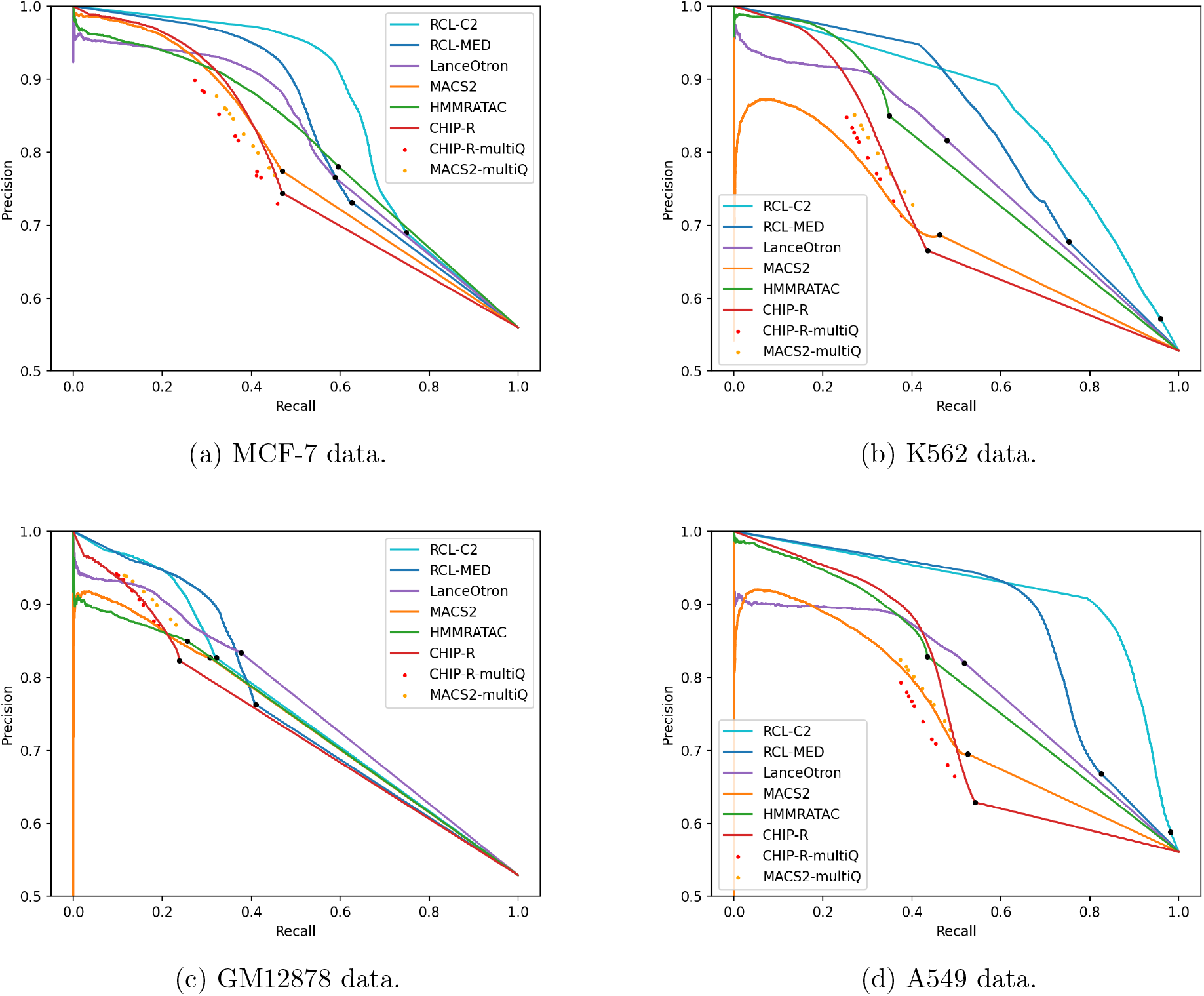
Precision-Recall (PR) curves for ChromHMM-labeled regions. Black dot in each curve denotes the region with lowest score; all remaining ChromHMM-labeled regions are not scored the method. RCL-C2, analysis with coverage threshold 2; RCL-MED, analysis with default “median” coverage threshold; MACSmultiQ and ChIP-R-multiQ dots are obtained by varying *q*-value cut-offs.

### 1.3 Performance benchmarking using transcription factor ChIP-seq data

In addition to genome annotations obtained with ChromHMM, we used transcription factor (TF) ChIP-seq data to evaluate method performance. These data mark potential binding sites of various TFs, which bind where DNA is accessible and should coincide with ATAC-seq peaks. RCL achieved the highest precision, except in the mouse placenta data where ChIP-R was more precise (Table 2c). As observed with ChromHMM labels, RCL precision improved upon lowering the threshold *t*.

### 1.4 Gene ontology analysis

As ChromHMM and TF ChIP-seq labels do not cover the whole genome and all methods predicted peaks outside these label regions, we analyzed the biological functions of genes associated with peaks called by each method (see §3.4 Gene ontology analysis in Methods). We expect meaningful peaks to associate with genes that are related to the known functions of the cell types or tissues. For example, we expect MCF-7 peaks to be enriched for processes such as epithelial cell proliferation, migration and invasion, as well as angiogenesis (Comşa et al. 2015). We therefore checked for the enrichment of any gene ontology (GO) term containing words “epithelial”, “epithelium”, or “angiogenesis”. The K562 cell line has antiapoptotic characteristics (Kuželová et al. 2004); therefore, we expect the enrichment of processes related to the negative regulation of apoptosis, and searched for terms that contained the words “apoptosis” or “apoptotic”. The cell line A549, a type of lung carcinoma epithelial cell, is an alveolar type II (ATII) cell that secretes surfactant protein to maintain homeostasis (Lee et al. 2018). Hence, processes underlying this cell type are related to terms that include “epithelial”, “epithelium” and “surfactant”. GM12878 is a human lymphoblastoid cell line generated by transforming primary B cells from peripheral blood with Epstein-Barr virus (EBV) (Bird et al. 1981; Anderson and Gusella 1984). Therefore, processes involving “B cell” should be enriched if biologically relevant peaks are supplied. Last, in the mouse placenta at day 9.5, the labyrinth layer is actively developing after chorioallantoic attachment finishes; as a result, a dense network of fetal blood vessels are forming within the layer where nutrients are exchanged (R. R. Starks et al. 2021; Cross et al. 2003; Watson and Cross 2005). In addition, the placenta is comprised mostly of trophoblast cells, which are epithelial-like cells. Thus, processes related to “placenta”, “epithelium”, “vasculature”, “angiogenesis”, “labybrinth” and “insulin” should be expected in meaningful peaks from day 9.5 mouse placenta tissue.

In general, we observed that only peaks uniquely called by RCL are enriched with relevant biological terms, with the exception of A549 data (Figure 3, Supplementary Figures S3–S6, Supplementary Tables S2–S6). For example, peaks that only RCL identified were associated with processes related to apoptosis in the K562 dataset (Figure 3). In case RCL benefitted from simply predicting a higher number of peaks, we randomly downsampled all peak sets and repeated the analysis. RCL continued to enrich on functionally relevant processes (Supplementary Figures S3–S6, Supplementary Tables S2–S6). For relevant terms, RCL peaks are often associated with at least five genes and have higher than two-fold enrichment (vertical line, Figure 3, Supplementary Figures S3–S6) unlike the other methods, suggesting they are more likely to be associated with relevant genes than peaks identified by competing methods. In summary, there is evidence that unique peaks predicted by RCL, not just those overlapping ChromHMM- or TF-derived labels, are biologically relevant.

**Figure 3:**
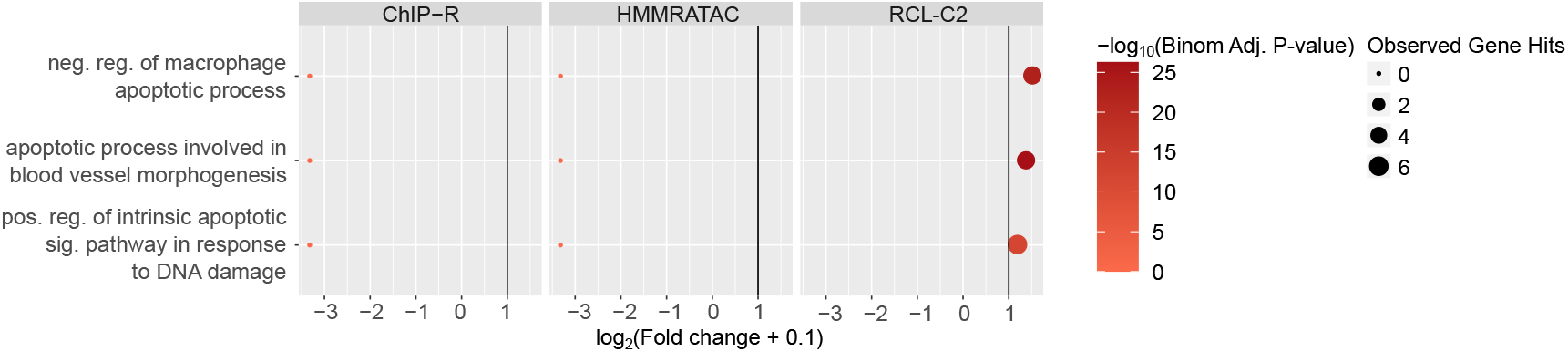
Gene ontology analysis using unique peaks called by each method in K562 data. Only relevant terms enriched with at least one peak set are plotted. Colors correspond to − log_10_(Binomial Adjusted P-value) where the adjustment was done following the Benjamini-Hochberg procedure (Benjamini and Hochberg 1995); dot sizes correspond to the observed number of genes associated with the term; *x*-axis corresponds to log_2_(Fold change+0.1) and vertical line is fold change of two. LanceOtron was not plotted since there was no unique peak called by the tool. Abbreviations: reg., regulation; pos., positive; neg., negative; sig., signaling.

## 2 Discussion

We propose RCL, an unsupervised peak caller for ATAC-seq data using contrastive learning across biological replicates. In our model, three losses: replicate similarity loss, class similarity loss and autoencoder loss, are learned simultaneously. We use ResNET as our backbone module with only five layers, making the network architecture shallow but efficient. On a server containing two Tesla V100 (16 GB) GPUs, the training time is 118 seconds when there are 4,828 1,000 bp regions and four replicates. Empirical results indicate RCL training time is roughly linear in the number of segments. In theory, training time is quadratic in the number of replicates because of the contrastive loss calculation, but replicate numbers remain quite low. Further investigation on timing is warranted, but total run times were acceptable on all datasets tested here. For example, for the A549 dataset (the largest dataset and slowest to train), the training time (25 epochs) took about 62 minutes.

In practice, only a small proportion of the genome is accessible (Consortium 2012). As a result, datasets for peak calling tend to be highly imbalanced, making it challenging to separate peak and non-peak regions. RCL showed no problems with class imbalance, probably because the region selection step effectively discards nonpeak regions and balances the data. If class imbalance proves to be a problem for calling datasets with sparser peaks or more widely across the genome in high coverage datasets, there are opportunities for improvement. For example, due to similarities to deep embedding clustering (Xie et al. 2016), cluster regularization methods proposed to avoid local optima or trivial solutions favoring predictions of the larger class (Zhong et al. 2020; Tao et al. 2018), may be applicable to contrastive learning and RCL.

Highly variable peaks or peaks in low coverage data may be difficult to find from single replicates, but their signal may become obvious when comparing across multiple replicates. HMMRATAC utilizes multiple replicates by combining them, which reduces the variance in the signal, but does not help the method learn what defines noise in a single replicate. ChIP-R, a post-hoc method to combine peaks called by another method, can improve performance over MACS, but only when used with a liberal *q*-value threshold followed by furthering filtering of ChIP-R-predicted peaks (red PR curves in Figure 2). Although both MACS and RCL make predictions for individual replicates, RCL predicts after learning from all replicates, while MACS predicts after learning from only the replicate in question. Currently, we combine the RCL prediction scores by taking the mean across replicates, but one can imagine more sophisticated approaches to combine predictions across replicates, possibly assessing the quality of prediction from each replicate and weighting the mean.

Replicates are, by design, an essential component of our method. To demonstrate the value of biological replicates we conducted an ablation study (Supplementary Text §S3.1.2). Contrasting real biological replicates gave the best predictions across chromosomes, which is not surprising given that biological replicates are fundamental for reproducibility and false signal reduction (F. Yan et al. 2020). In the absence of biological replicates, contrasting with an augmentation of the available data is better than contrasting with self. It could be that noise along the genome recapitulates some of the noise between biological replicates, but more study is necessary to understand RCL performance in the absence of replicates. Experiments varying the number of replicates available to RCL showed little effect on performance, even when the added replicate had substantially higher coverage (Supplementary Text §S3.3). All the experimental data examined in the current study have used high quality data with minimal batch effects and samples mostly taken from cultured cells with likely little biological variation, all of which may explain the limited impact of additional replicates. It will be an interesting future direction to examine how contrastive learning and the RCL framework handle batch effects or the inclusion of low quality replicates.

We acknowledge that the labeled regions indicating the “ground truth” used for assessment are noisy. First, the annotations obtained using ChromHMM (Ernst and Kellis 2017) applied to several ChIP-seq datasets and thus innately contain technical noise from data generation and model estimation. The TF ChIP-seq labels were specifically called by MACS2 (Zhang et al. 2008), which we know produces noisy, imperfect labels. Second, while we matched cell types and biological conditions, variation in the samples used to generate TF ChIP-seq or ChromHMM labels were not completely controlled. Notably in the mouse placenta data, the biological condition did not exactly match between the TF ChIP-seq and ATAC-seq data, so this set of labels contains additional biological noise. Third, our translations from ChromHMM states to open/closed regions were imperfectly determined to the best of our knowledge. There appear to be noisy truth labels in the MCF-7 data. Some negative ChromHMM regions were assigned high scores (logit-transformed scores *>* 10) (Supplementary Figure S9a; see Supplementary Figure S9 for a study of scores from all methods and datasets). Although these score assignments could be due to the shortcomings of RCL, it is also possible that some labels are wrong. Similar issues likely arise in the labels provided to supervised methods (Oh et al. 2020; Hentges et al. 2021), which is another reason to further develop unsupervised peak calling.

When there is noise in the labels, the observed performance metrics (precision, recall, F1, and PR curve) are not equal to the true performance metrics evaluated against the truth (Jiang et al. 2014). Most worryingly, the observed recall is a function of true recall *and* the true false positive rate. Specifically, let *ŷ* be predicted labels, *y* unobserved true labels, and *z* observed noisy labels. Further, suppose the labeling error rates *P* (*z* = 0 | *y* = 1) = *P* (*z* = 1 | *y* = 0) = *E* are constant and independent of any signal in the data. Then, the observed recall is

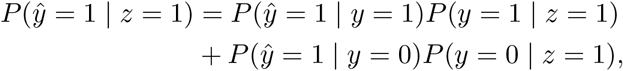

where *P* (*ŷ* = 1 | *y* = 0) is the true false positive rate (FPR). Thus, observed recall is a contaminated measure of recall, and methods compared via observed recall (or F1 or PR curve) may not reveal their actual ranking. Given this concern, it is possible to estimate method performance *in the context of label errors* (Raykar et al. 2009; Y. Yan et al. 2014) or correct errored labels so traditional assessment metrics are more accurate (Sabetpour et al. 2021; G. Zheng et al. 2021). Alternatively, performance evaluation can be carried out with simulated data. However, there is no existing simulation method for ATAC-seq data, and the existing methods used for ChIP-seq, such as A. Zheng et al. (2022), are not applicable for ATAC-seq.

RCL learns and predicts on fixed sized segments (length *α*, default 1,000 bp). We did not examine the impact of hyperparameter *α* on RCL performance, but it certainly complicates peak calling. We chose to transfer RCL prediction scores from the *α* bp segments to the variable-length candidate regions produced by the algorithm of §1.1.1. Prediction region selection, because it works well (Supplementary Text §S3.4). Using these candidate regions with mean coverage as a simple score already does well in MCF-7, but RCL learns additional signals, perhaps peak shape, that further improve the performance (Supplementary Figure S11a). Not only do the candidate regions work well, but they are not easily substituted.

Using the *α* bp segments as peak predictions in A549 failed, probably because they lack the resolution to pinpoint narrow peaks, but a quick and dirty attempt to shrink the prediction regions to the relevant peak summit performed even worse (Figure S11b). A better solution may be to learn and predict directly on the variable-sized candidate regions. We could pad variable-sized inputs to the same length or we could add a Spatial Pyramid Pooling layer (He et al. 2015) before the first fully-connected layer to remove the fixed-size constraint of the network. On the other hand, such an approach would still require data preprocessing to choose the candidate regions. An even better solution might be to predict at the nucleotide level, a one-step solution to identify peaks and their extent.

RCL can be extended and improved in other ways. First, we used simple read coverage as input, but HMMRATAC reports reproducible signal in the coverage of distinct fragment lengths around open regions (Tarbell and Liu 2019). RCL could be extended to take coverage vectors for multiple fragment lengths, the fragments themselves, or even annotation information, as used by the supervised method CNN-Peaks (Oh et al. 2020). Second, multiple hyperparameters in both data processing and model training can be further tuned. For example in input preparation, regions longer than 100bp are kept in the current method. We have tried keeping regions longer than 147bp, and it resulted in fewer inputs and fewer called peaks; however, we still obtained good predictions. Last, we have focused on ATAC-seq data, where peak calling has been particularly difficult because of the lack of control samples and good truth labels. Nevertheless, our model assumes nothing particular to ATAC-seq data and can be applied to ChIP-seq, CUT&RUN (Skene et al. 2018) and other techniques requiring peak calling.

There is clearly much left to learn about how RCL works to extract useful signal from replicates, but we can offer some preliminary recommendations. First, we recommend users follow established data preprocessing and quality control steps for ATAC-seq data (F. Yan et al. 2020). Where we have tested, few hyperparameters and inputs of RCL had much impact on performance, other than the coverage threshold *t*, option `-t`, and the candidate regions. The default threshold (“median”) identified highly confident peaks with excellent precision (Figure 2); therefore, this setting can be a good starting point for researchers to find the highest confident peaks. If a researcher wishes to predict more peaks accurately, it may be better to reduce threshold *t* and expose RCL to more and less obvious candidate peaks. This is a particularly good option for high coverage datasets, where RCL reproducibly outshines the competing methods. We recommend using all replicates under the assumption that replicates are still quite sparse because of cost. While additional replicates did not improve performance on datasets tested here, they also did not hurt performance. Finally, we provide no options to explore alternative prediction regions, but users may have good ideas for choosing candidate regions and they can try them out with the RCL software package.

In summary, we have developed a novel peak calling framework for ATAC-seq data using contrastive learning techniques to extract signals shared across biological replicates and identify high confident open chromatin regions. Because RCL can predict more peaks with higher precision, it will facilitate future epigenome and chromatin accessibility studies in various biological contexts.

## 3 Methods

### 3.1 ATAC-seq data acquisition

ATAC-seq data sets of the following human cell lines and mouse tissues were obtained from public databases: MCF-7, A549, K562, GM12878, and mouse placenta. The MCF-7 dataset, with two biological replicates, was accessed through the ENCODE experiment ID ENCSR422SUG (Consortium 2012). The A549 dataset, with three biological replicates, was accessed through the ENCODE experiment ID ENCSR032RGS (Consortium 2012). The K562 dataset, with three biological replicates, was accessed through the ENCODE experiment ID ENCSR868FGK (Consortium 2012). The GM12878 dataset generated using 50, 000 cells was obtained from four replicates with the accession numbers SRR891268, SRR891269, SRR891270 and SRR891271 (Buenrostro et al. 2013). Last, the mouse data generated from mouse placenta at day 9.5, with three biological replicates, was accessed with the accession numbers SRR7912013, SRR7912014 and SRR7912015 (R. Starks et al. 2019).

### 3.2 ATAC-seq data processing

FASTQ files were assessed using FastQC (Andrews 2010) (version 0.11.7) to identify samples with over-represented sequences or adapter contamination. Trimmomatic (Bolger and Giorgi 2014) was used to remove adapter content and filter low quality base pairs and reads (`ILLUMINACLIP:overpresentedSeq.fa:2:30: 10:2: keepBothReads LEADING:3 TRAILING: MINLEN:36`, other settings: default, version 0.39). Here, the `overpresentedSeq.fa` file contains the over-represented sequences and adapter content identified with FastQC. Reads were aligned to the autosomal and sex chromosomes of human reference genome GRCh38 or mouse reference genome GRCm38 (release 98) (Cunningham et al. 2019) using Bowtie2 (Langmead and Salzberg 2012) (`-X 1000 --no-discordant`, other settings: default, version 2.3.4.1). Picard (Broad Institute 2019) was used to remove duplicate reads (`REMOVE_DUPLICATES=true`, version 2.17.0). Reads with low quality mapping (`MapQ<20`) were removed before merging, sorting, and indexing the resulting BAM files with SAMtools (Danecek et al. 2021). Last, to assess sample quality after preprocessing, ataqv (Orchard et al. 2020) (`--ignore-read-groups`, other settings: default, version 1.2.1) was used to check for fragment length distribution and transcription start site (TSS) enrichment. Samples used for downstream analyses must have a mononucleosome peak in the fragment length distribution, and TSS enrichment ≥ 1.5.

### 3.3 Tuning RCL

We used dilation 8 and kernel size 31 to train our model. Other hyperparameters are set to be default values (number of epochs = 25, batch size = 256, learning rate = 10^−4^ and temperature *τ*_1_ = *τ*_2_ = 0.5). Details regarding choosing dilation 8, kernel size 31 and model development were are discussed in Supplementary Text §S3.1. Briefly, RCL was developed on the MCF-7 cell line data (see §3.1. ATAC-seq data acquisition) using different truth labels than those presented in results. We will demonstrate that a *roughly* tuned RCL is already substantially superior to existing methods, not only on MCF-7 with a distinct truth, but on additional holdout datasets as well.

### 3.4 Method comparison

We compared RCL to MACS (Zhang et al. 2008), ChIP-R (Newell et al. 2021), HMMRATAC (Tarbell and Liu 2019), and LanceOtron (Hentges et al. 2021). Call performance was assessed using three analyses: comparisons using truth labels of genome annotation obtained with ChromHMM (Ernst and Kellis 2017) from independent data collected on the same cell lines and tissues; comparisons using truth labels of transcription factor ChIP-seq data collected on the same cell lines and tissues; and the association of peak prediction to biologically relevant genes.

#### MACS peak calling

MACS (version 2.1.1) (Zhang et al. 2008; Gaspar 2018) was used to call peaks with BAM files from individual replicates. Peaks were called with options `-g hg -f BAMPE --bdg --keep-dup all`, with the following cut-offs for the *q*-value `-q: 0.5, 0.1, 0.05, 0.01, 0.005, 0.001`, `0.0005, 0.0001, 0.00002`, and `0.00001`, and other settings: default. Any peaks overlapping with a blacklist region (Amemiya et al. 2019) by at least 1 bp are removed. MACS was originally developed for calling peaks on transcription factor ChIP-seq data, so the default settings and model assumptions may not apply for ATAC-seq data. We have some evidence that altering shift and window sizes can improve MACS performance in some aspects (Supplementary Text S3.5), but settings to consistently improve MACS performance were elusive and beyond the scope of this work. Given peak calls from individual replicates, the peak union method was used to combine peaks across replicates. Specifically, a consensus peak set is the union of peaks overlapping with each other by ≥ 50% length in ≥ 2 replicates. Scores of consensus peaks were the mean − log_10_(*q*-value) at peak summit of the individual peaks observed in separate replicates. As MACS does not report scores of non-peak regions, replicates not calling a peak in the region are not used when calculating scores of consensus peaks.

#### ChIP-R peak calling

We used ChIP-R (version 1.1.0) (Newell et al. 2021) as an additional, independent method for combining peaks called from MACS. Peaks were first called with MACS as described above. Then, ChIP-R was run with the following setting: `-m 2`, `-a 0.5, 0.1, 0.05, 0.01, 0.005, 0.001, 0.0005, 0.0001,0.00002,` and `0.00001` (matching with `-q` in MACS), other settings: default. Any peaks overlapping with a blacklist region (Amemiya et al. 2019) by at least 1 bp are removed. The reported `score` was used as scores of ChIP-R peaks.

#### HMMRATAC peak calling

HMMRATAC (version 1.2.4) (Tarbell and Liu 2019) was used to call peaks with a merged BAM file from all replicates, options `-Xmx128G`, `--window 250000`, other settings: default. A peak is a region in the open state with scores ≥ 0, reported by default with the `--peaks` option. By default, peak scores of HMMRATAC are the maximum read coverage of the called center state region. Any peaks overlapping with a blacklist region (Amemiya et al. 2019) by at least 1 bp were removed.

#### LanceOtron peak calling

To implement et al. 2021) (ver- (Hentges LanceOtron sion 1.0.8), the input bigwig files were obtained using deeptools (version 2.5.2) (Ramírez et al. 2016) with the following command: `bamCoverage -b bamFile -o bigwigFile --extendReads -bs 1 --normalizeUsingRPKM`. The inputs were then used to call peaks with default settings and LanceOtron’s pretrained model. Any resulting regions overlapping with a blacklist region (Amemiya et al. 2019) by at least 1 bp are removed. Then, the union of regions overlapping with each other by ≥ 50% length in ≥ two replicates were obtained as a consensus region set. Scores of consensus regions were the mean `overall_peak_score` of the individual regions. Regions with scores ≥ 0.5 were then defined as peaks by default.

#### RCL peak calling

RCL was used with coverage threshold *t* of “median” and 2, other settings: default. By default, any segment overlaps with blacklist regions was excluded due to the segment selection procedure. Let peak prediction score given by RCL be *ξ*_*ri*_ = *q*_*ri*1_ for the *i*th *α* bp segment in the *r*th replicate. We obtain a final peak prediction score for each region in 𝒜 (see Step 2 in §1.1.1. Prediction region selection) by averaging over *ξ*_*ri*_ for all replicates *r* = 1, 2, …, *R* and segments *i* extracted from the region. A region in 𝒜 is predicted to contain at least one peak if this score is ≥ 0.5.

#### Compilation of true positive and true negative labeled regions by ChromHMM

For human cell line data, we obtained genome annotations inferred with ChromHMM (Ernst and Kellis 2017) from ENCODE. Specifically, genome annotation for MCF-7 data was accessed via the experiment ID ENCSR579CCH, the A549 data via ENCSR283FYU, and the GM12878 data via ENCSR988QYW. True positive regions are those marked “EnhA1”, “EnhA2”, “EnhG1”, “EnhG2”, “TssA”, “TssFlnk”, “TssFlnkD”, “TssFlnkU”, and “Tx”. True negative regions are marked as “Het”, “Quies”, “ReprPC”, and “ZNF/Rpts”. Annotations not in these lists were not used, and regions overlapping with a blacklist region (Amemiya et al. 2019) by at least 1 bp were removed. For full definitions of the states, see Supplementary Table S7.

For the mouse placenta data, we obtained ChromHMM annotation from (R. R. Starks et al. 2021). True positive regions are those belonging to State 8, 9 and 10, and true negative regions are those belonging to State 2. Detailed biological characterization of these states were described in (R. R. Starks et al. 2021).

#### Compilation of true positives from transcription factor (TF) ChIP-seq data

For human cell line data, we obtained TF ChIP-seq data from matching cell lines from ENCODE (Consortium 2012). Bed files of IDR thresholded peaks were downloaded from all datasets that passed all quality control criteria of ENCODE and had “released” status, and their bio-samples were not perturbed. For mouse placenta data, no TF ChIP-seq data from matching condition was available. Therefore, we obtained TF ChIP-seq data from mouse differentiated trophoblast stem cells. GEO accession IDs of datasets were searched through the GTRD database (Yevshin et al. 2018) with species “Mus musculus”, type “ChIP-seq”, cells “differentiated trophoblast stem cells”, and the experiments were not subjected to any special treatments (treatments were “Wild type” or “None”). The peak files were obtained from their GEO pages. If the data was originally aligned to mm9, the peaks were moved to mm10 coordinates with the LiftOver tool (Hinrichs et al. 2006) (default settings).

True positive regions were defined as those with at least one TF ChIP-seq peak. Regions overlapping with a blacklist region (Amemiya et al. 2019) by at least 1 bp were removed. No true negative regions were defined using these datasets. For lists of data used, see Supplementary Table S7.

#### Calculation of evaluation metrics

Since labelled regions and called regions do not necessarily coincide, we defined a mapping function to transfer scores of called regions in the ATAC-seq data to predicted scores for annotated regions. Specifically, suppose there are *n*_*i*_ called regions overlapping with the *i*th labeled region, and *c*_*j*_ (1 ≤ *j* ≤ *n*_*i*_) is the predicted score that overlaps by *o*_*j*_ base pairs with the *i*th labeled region. Then the weighted prediction score for the *i*th labeled region is 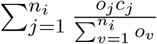. In case *n*_*i*_ = 0, we assign the lowest weighted score observed for that method as the predicted score.

We used point estimates of precision, recall and F1, as well as Precision-Recall (PR) curves to compare the performance of the methods. Specifically, for MACS and ChIP-R, we calculated precision and recall for each `-q` (`-a`) cut-off. For plotting PR curves, we used MACS or ChIP-R with *q*-value 0.5. Results from other *q*-value settings were presented as point estimates on the plots. RCL can call more peaks by lowering the threshold *t*, but changing *t* will also change the fitted model and the peak calls made. We always ran RCL with defaults, but we also selected non-default *t* so the *number* of candidate peaks roughly matched the number of peaks called by MACS (*q*-value 0.5), ChIP-R (*q*-value 0.5) and HMMRATAC.

#### Gene ontology analysis

To assess the potential functional roles of the called peaks by all methods, we used gene ontology analysis. We examined the following sets of peaks. For the MCF-7, K562 and A549 data, we used ChIP-R, HMMRATAC, LanceOtron, and RCL peaks called when *t* = 2 peaks; for the GM12878 and mouse placenta data, ChIP-R, HMMRATAC, LanceOtron, and RCL peaks called when *t* = *median* peaks; and for all datasets, peaks identified uniquely by each of these methods. Specifically, a peak is uniquely assigned to a method if it does not overlap with peaks predicted by any other method, as assessed using BEDTools `intersect -v` (Quinlan and Hall 2010).

We used the Genomic Regions Enrichment of Annotations Tool (GREAT) (version 4.0.4) (McLean et al. 2010; Tanigawa et al. 2022) implemented in R (Gu and Hübschmann 2022) to carry out GO enrichment using either the human GRCh38 or mouse GRCm38 annotations and the default basal plus extension association rule. For each analysis, we randomly selected peaks so that the number of input regions for GREAT was the smallest or second smallest peak set size amongst all tools (see number of peaks from each tool in Supplementary Table S1). For unique peaks, we also analyzed all peaks without down-sampling. A biological process term was considered enriched if its binomial *q*-value ≤ 0.05, binomial fold change ≥ 2, and the observed number of associated genes was ≥ 5.

## 4 Data Accessibility

All data used in the study were listed in the Methods section and Supplementary Table S7.

## 6 Acknowledgements

We acknowledge the Research IT group at Iowa State University (http://researchit.las.iastate.edu) for providing servers and IT support. We would like to thank Dorman lab and Tuteja lab members for their discussion and support. This work was supported in part by the Eunice Kennedy Shriver National Institute of Child Health & Human Development of the National Institutes of Health under award number R01HD096083 (to G Tuteja). G Tuteja is Pew Scholar in the Biomedical Sciences, supported by The Pew Charitable Trusts. This work was supported in part by the United States Department of Agriculture (USDA) National Institute of Food and Agriculture (NIFA) Hatch project IOW03717. The findings and conclusions in this publication are those of the author(s) and should not be construed to represent any official USDA or U.S. Government determination or policy.

