## Supplementary materials for "Unsupervised Contrastive Peak Caller for ATAC-seq"

#### S1 Supplementary Figures

This section includes supplementary figures cited to support claims in the main text. Figure S1 displays the distribution of threshold values used in extracting candidate regions for RCL training. Figure S2 plots the PR curves for the mouse day 9.5 placenta data, which was excluded from main text Figure 2 for space. Figures S3–S6 show a more detailed Gene Ontology enrichment analysis accompanying Results §1.4.

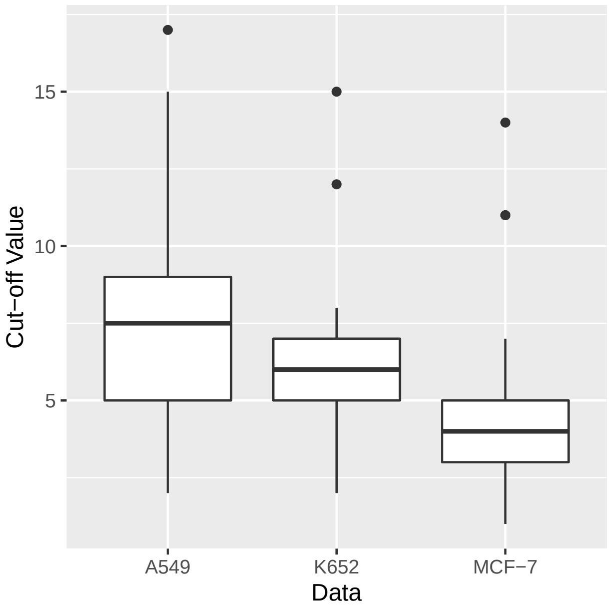


Figure S1: Distribution of default threshold (cut-off) values for extracting candidate peak regions (see main text §1.1.1) across chromosomes. The GM12878 and day 9.5 mouse placenta datasets are not shown since their median coverage (among all covered sites) is one for all chromosomes.

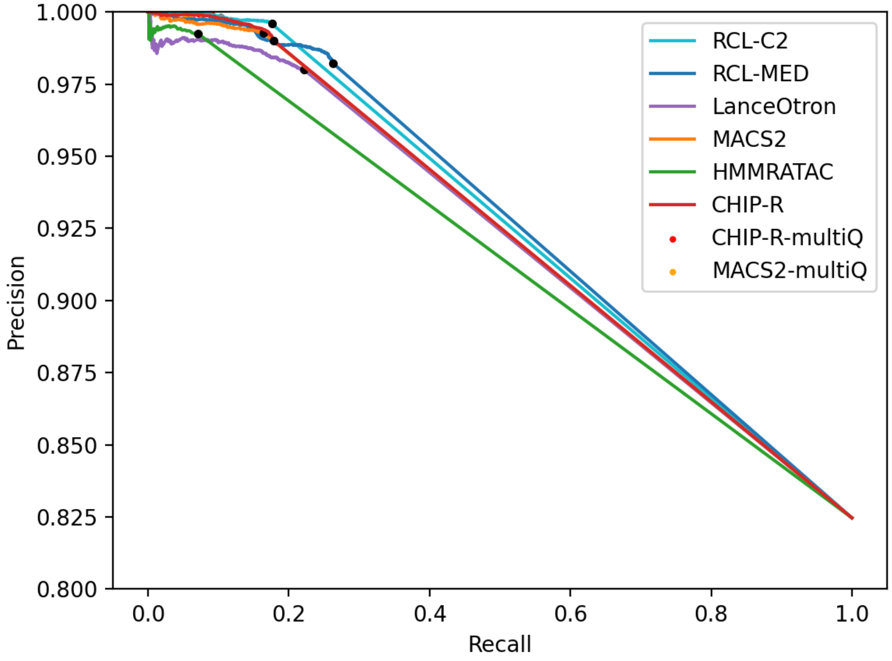


Figure S2: Precision-Recall (PR) curve for mouse placenta data. See main text Figure 2 for PR curves for other datasets and more information. ChIP-R and MACS2 multiQ points at the upper, left have been overplotted by PR curves.

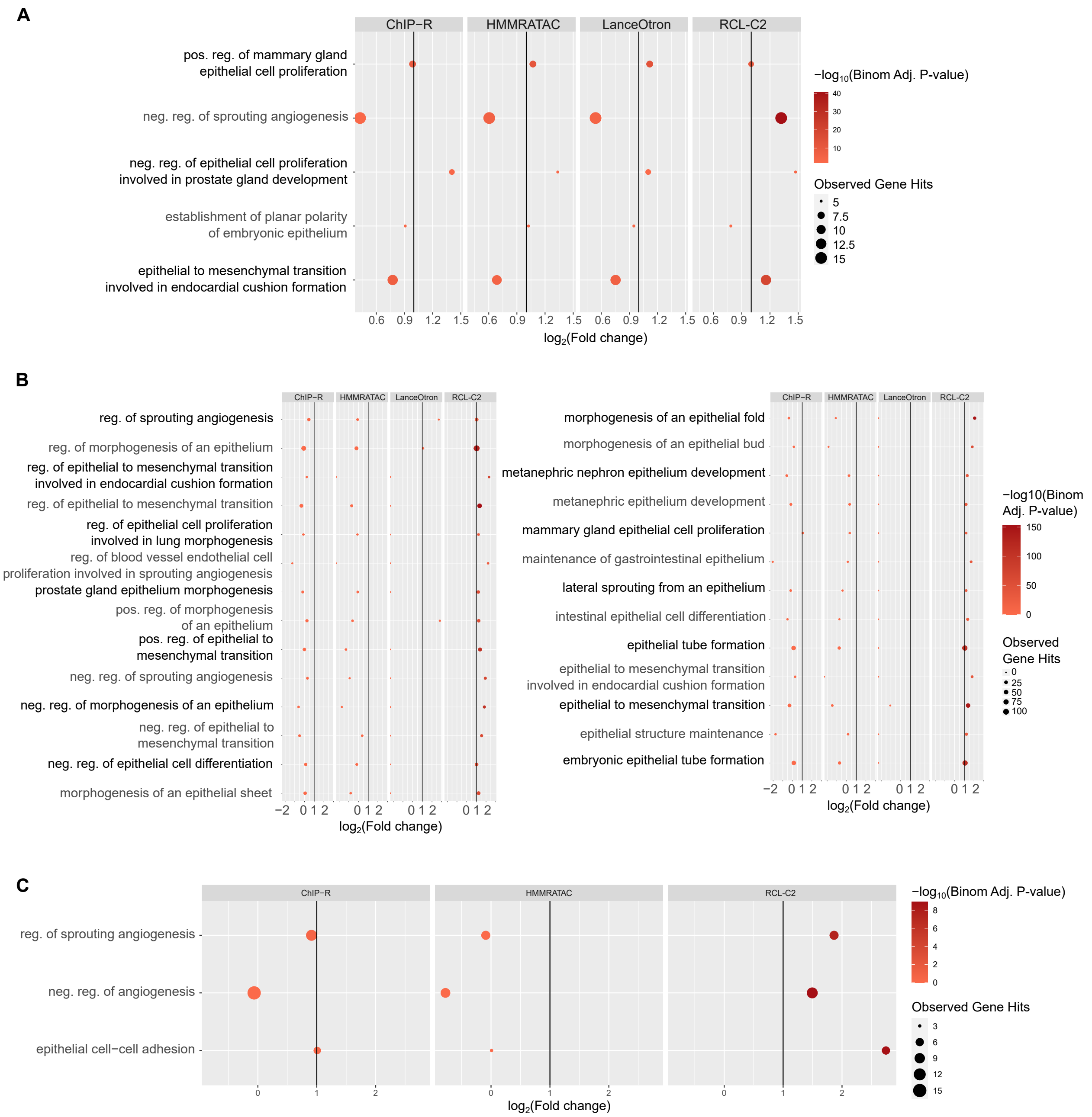


Figure S3: Gene ontology analysis of MCF-7 data. Peaks included in analysis are (A) randomly downsampled from all peaks to the smallest number of peaks found by any method; (B) all unique peaks found by each tool; (C) randomly downsampled from unique peaks to the second smallest number of unique peaks found by any method. LanceOtron unique peaks were not included in (C) due to its small number of peaks (see Supplementary Table S1). Only relevant terms enriched in at least one peak set are plotted. Colors correspond to $-\log_{10}$(Binomial Adjusted P-value) where the adjustment was done following the Benjamini-Hochberg (Benjamini and Hochberg 1995) procedure; dot sizes correspond to the observed number of genes associated with the term; $x$-axis corresponds to $\log_2$(Fold change) and vertical line corresponds to fold change of two, which was used to indicate biological significance. Abbreviations: reg., regulation; pos., positive; neg., negative; sig., signaling.

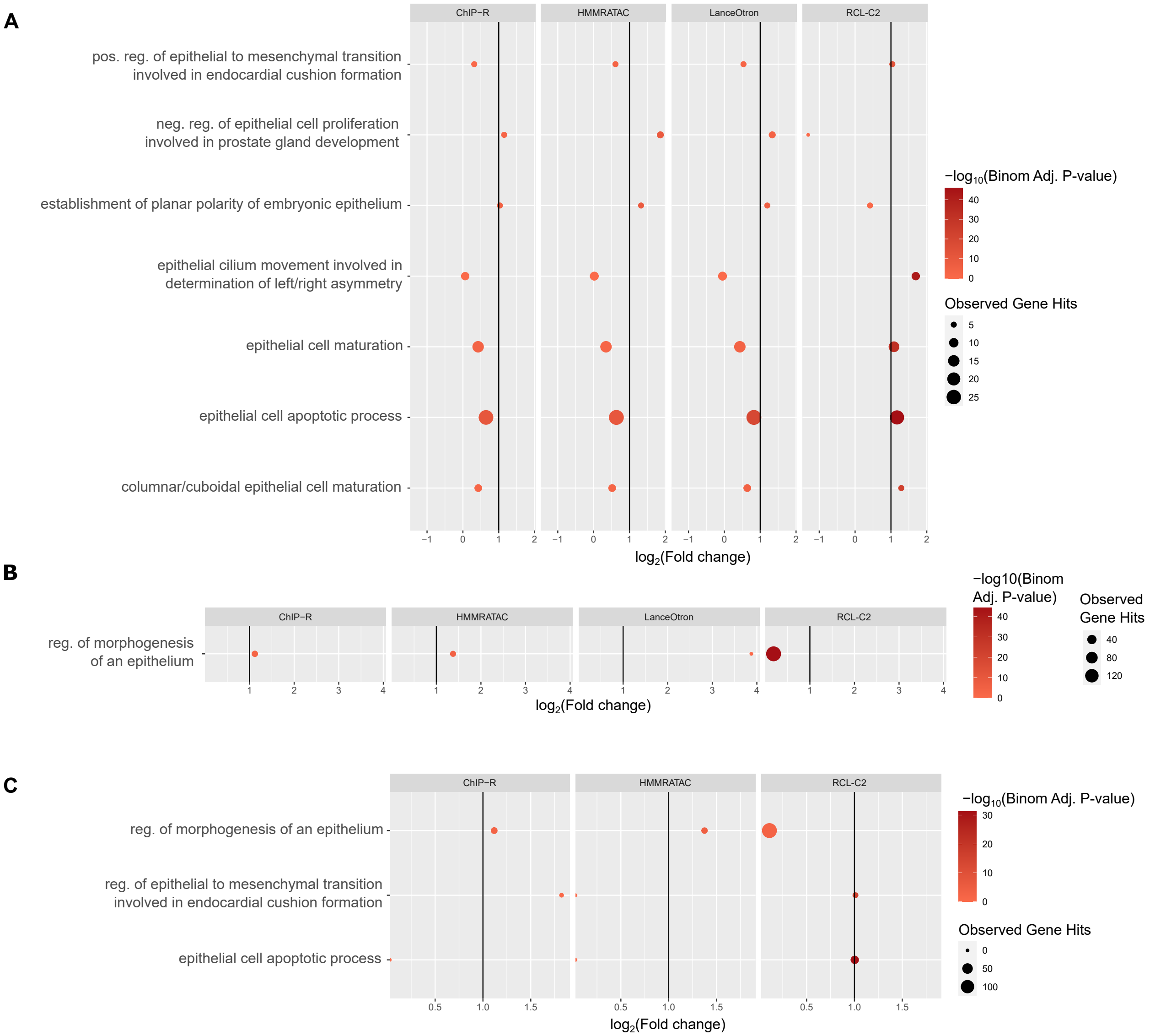


Figure S4: Gene ontology analysis of A549 data. Peaks included in analysis are (A) randomly downsampled from all peaks to the smallest number of peaks found by any method; (B) all unique peaks found by each tool; (C) randomly downsampled from unique peaks to the second smallest number of unique peaks found by any method. LanceOtron unique peaks were not included in (C) due to its small number of peaks (see Supplementary Table S1). Only relevant terms enriched in at least one peak set are plotted. See Figure S3 legend for details.

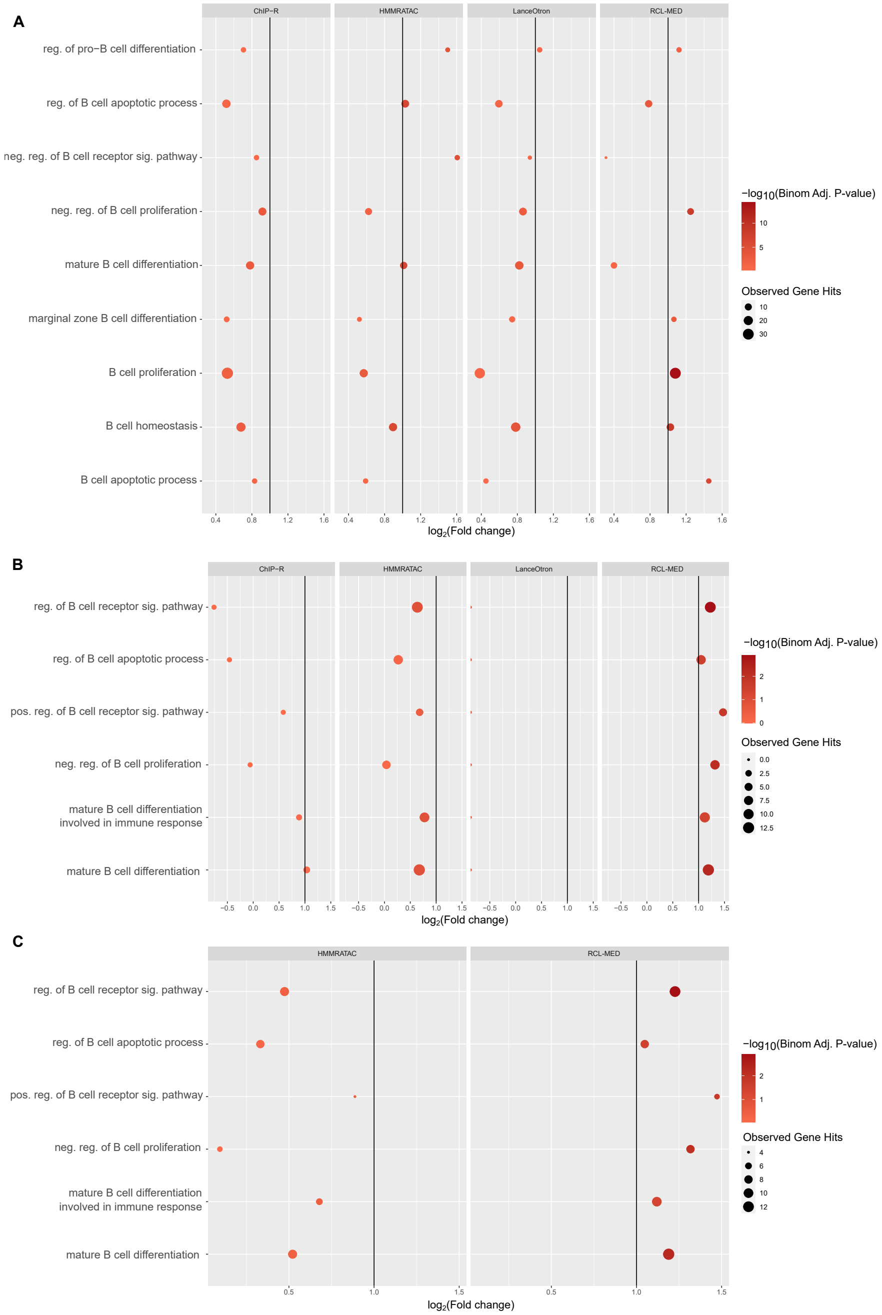


Figure S5: Gene ontology analysis of GM12878 data. Peaks included in analysis are (A) randomly downsampled from all peaks to the smallest number of peaks found by any method; (B) all unique peaks found by each tool; (C) randomly downsampled from unique peaks to the third smallest number of unique peaks found by any method. ChIP-R and LanceOtron unique peaks were not included in (C) due to their small numbers of peaks (see Supplementary Table S1). Only relevant terms enriched in at least one peak set are plotted. See Figure S3 legend for details.

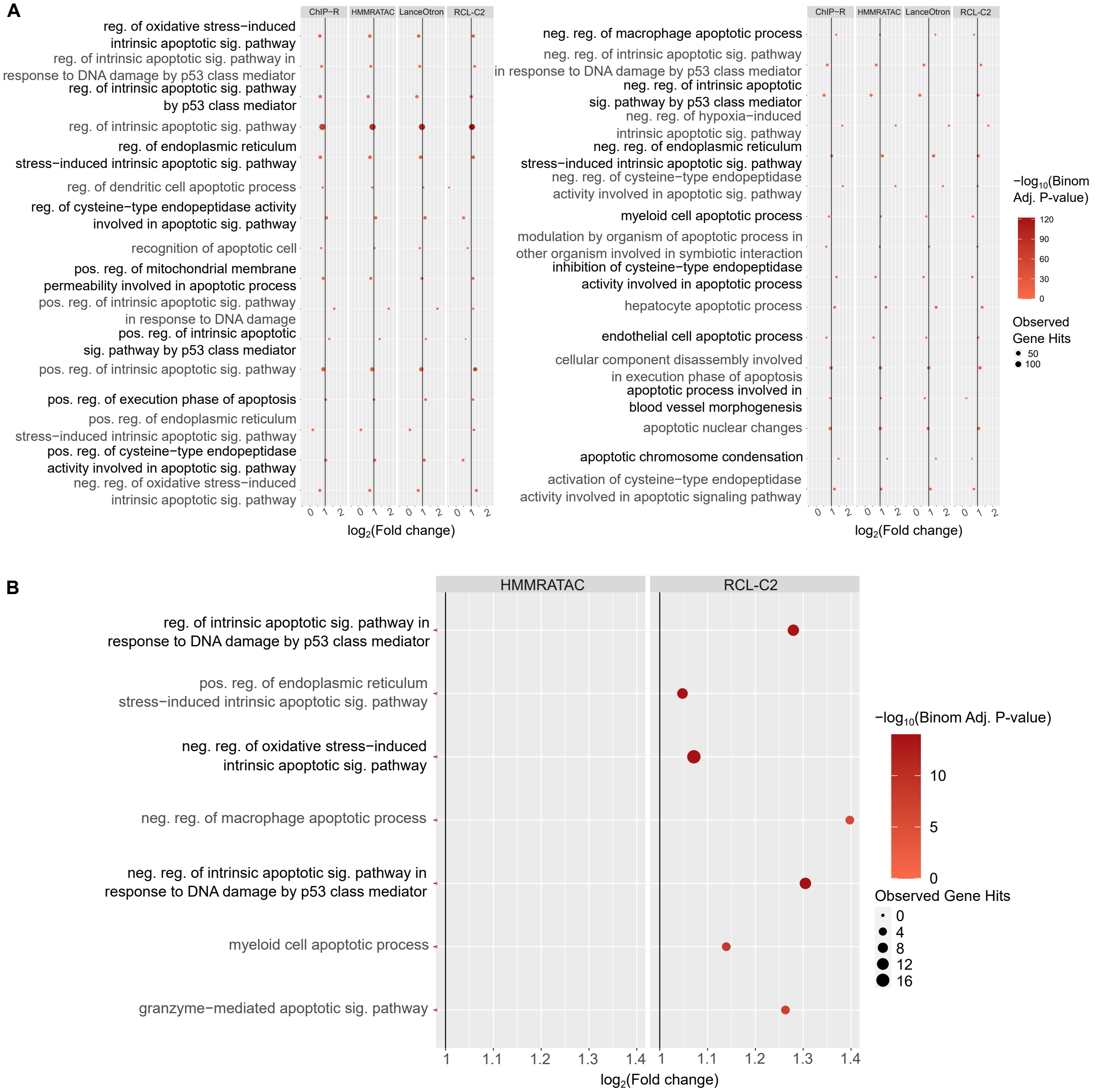


Figure S6: Gene ontology analysis of K562 data. Peaks included in analysis are (A) randomly downsampled from all peaks to the smallest number of peaks found by any method; (B) all unique peaks found by each tool; (C) randomly downsampled from unique peaks to the third smallest number of unique peaks found by any method. ChIP-R and LanceOtron unique peaks were not included in (C) due to their small numbers of peaks (see Supplementary Table S1). Only relevant terms enriched in at least one peak set are plotted. Abbreviations: reg., regulation; pos., positive; neg., negative; sig., signaling.

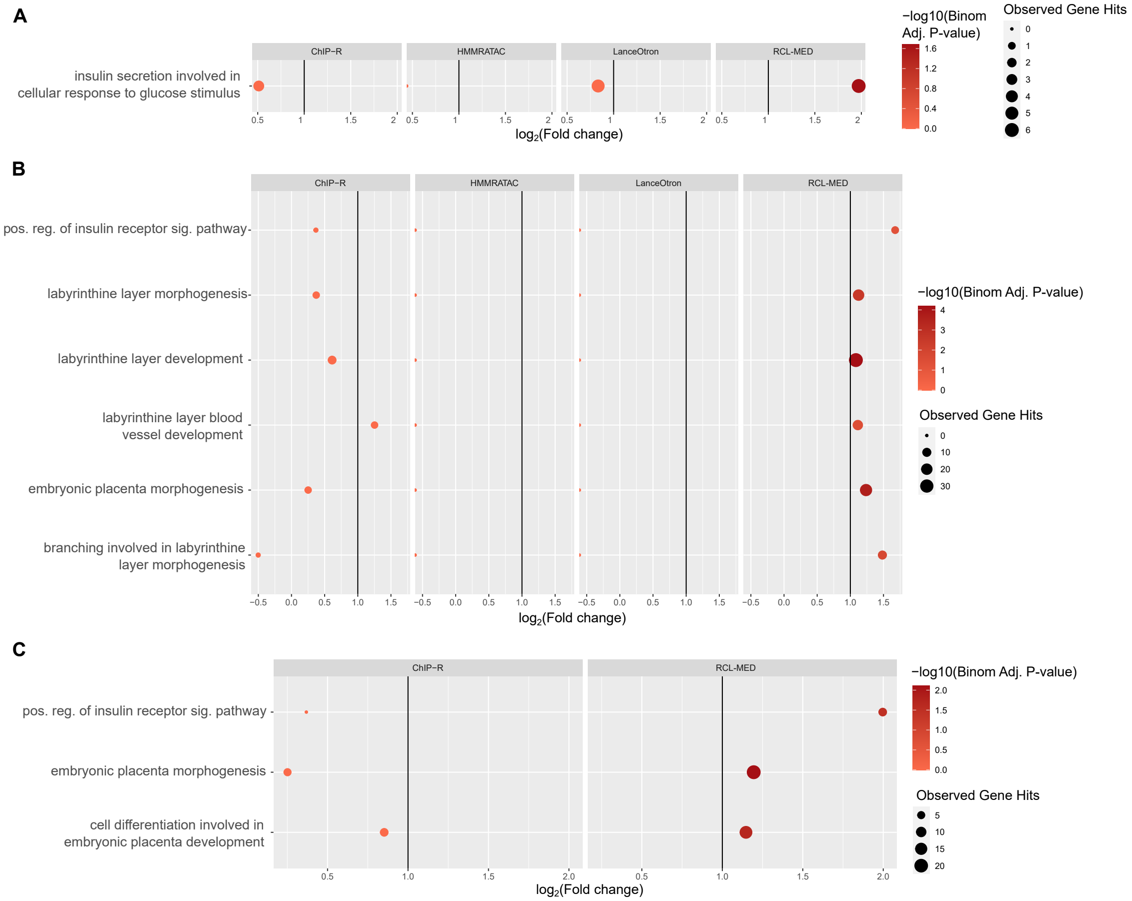


Figure S7: Gene ontology analysis of day 9.5 mouse placenta data. Peaks included in analysis are (A) randomly downsampled from all peaks to the smallest number of peaks found by any method; (B) all unique peaks found by each tool; (C) randomly downsampled from unique peaks to the third smallest number of unique peaks found by any method. HMMRATAC and LanceOtron unique peaks were not included in (C) due to their small numbers of peaks (see Supplementary Table S1). Only relevant terms enriched in at least one peak set are plotted. Abbreviations: reg., regulation; pos., positive; neg., negative; sig., signaling.

#### S2 Supplementary Tables

Supplementary Table S1 reports the number of peaks called by each method, including unique peaks not overlapping with peaks called by any other method. Table 1 reports the number of called peaks overlapping with ChromHMM labels. All supplementary tables, including S1, are also available as excel spreadsheets.

| | | Number of Peaks | |
|---|---|---|---|
| **Dataset** | **Method** | **Total** | **Unique** |
| MCF-7 | MACS2 ($q = 0.05$) | 68,694 | NC |
| | ChIP-R ($q = 0.05$) | 69,864 | 5,201 |
| | MACS2 ($q = 0.5$) | 73,817 | NC |
| | ChIP-R ($q = 0.5$) | 99,022 | NC |
| | HMMRATAC | 69,885 | 1,732 |
| | LanceOtron | 54,771 | 187 |
| | RCL-MED | 58,057 | NC |
| | RCL-C2 | 108,886 | 32,005 |
| A549 | MACS2 ($q = 0.05$) | 94,659 | NC |
| | ChIP-R ($q = 0.05$) | 173,664 | 650 |
| | MACS2 ($q = 0.5$) | 130,063 | NC |
| | ChIP-R ($q = 0.5$) | 304,311 | NC |
| | HMMRATAC | 69,191 | 595 |
| | LanceOtron | 68,442 | 3 |
| | RCL-MED | 205,518 | NC |
| | RCL-C2 | 603,441 | 479,679 |
| GM12878 | MACS2 ($q = 0.05$) | 30,485 | NC |
| | ChIP-R ($q = 0.05$) | 28,399 | 654 |
| | MACS2 ($q = 0.5$) | 48,333 | NC |
| | ChIP-R ($q = 0.5$) | 57,234 | NC |
| | HMMRATAC | 44,664 | 6,718 |
| | LanceOtron | 17,943 | 2 |
| | RCL-MED | 47,060 | 5,289 |
| | RCL-C2 | 28,628 | NC |
| K562 | MACS2 ($q = 0.05$) | 80,554 | NC |
| | ChIP-R ($q = 0.05$) | 92,036 | 77 |
| | MACS2 ($q = 0.5$) | 119,547 | NC |
| | ChIP-R ($q = 0.5$) | 144,877 | NC |
| | HMMRATAC | 61,293 | 297 |
| | LanceOtron | 63,817 | 0 |
| | RCL-MED | 163,852 | NC |
| | RCL-C2 | 297,469 | 213,680 |
| Mouse placenta | MACS2 ($q = 0.05$) | 7,047 | NC |
| | ChIP-R ($q = 0.05$) | 9,797 | 1,092 |
| | MACS2 ($q = 0.5$) | 17,666 | NC |
| | ChIP-R ($q = 0.5$) | 30,787 | NC |
| | HMMRATAC | 229 | 27 |
| | LanceOtron | 3,179 | 9 |
| | RCL-MED | 12,873 | 4,412 |
| | RCL-C2 | 5,621 | NC |

Table S1: Number of called peaks and uniquely called peaks by each method for each dataset. A uniquely called peak is one that does not overlap with a peak called by any other method. NC: not calculated.

### S3 Supplementary Text

#### S3.1 Model Development

Data from the MCF-7 cell line (see Methods section 3.1) was used to develop the RCL method. During this process, we used different truth labels and procedures for calling peaks than those used in final method assessment. In particular, we used RCL to identify candidate peaks as those $\alpha$ bp segments $i$ with average peak probability $\sum_{r=1}^{R} q_{ri1}/R > 0.5$, where $q_{ri1}$ is the probability segment $i$ is a peak in replicate $r$ as defined in Section 1.1. Then, we defined candidate peak segments to be true open segments if they overlapped ($\geq$ 1 bp) regions marked by H3K27ac histone modification in relevant ChIP-seq data (peaks from ENCODE experiment ID ENCFF491LQY, called by MACS2). This histone modification was chosen since it marks active promoters and active enhancers across the genome (Creyghton et al. 2010). Here, only autosomal chromosomes were used. The H3K27ac histone mark was just one of at least six marks available to ChromHMM during annotation (Kundaje et al. 2015), so the labels used in RCL model development are related but not identical to the labels used in method assessment.

##### S3.1.1 RCL hyperparameter tuning

RCL involves several hyperparameters. The backbone model is composed of multiple CNN layers, with dilation and kernel size hyperparameters that we wanted to optimize. Other hyperparameters are set to standard default values (number of epochs = 25, batch size = 256, learning rate = $10^{-4}$ and temperature $\tau_1 = \tau_2 = 0.5$). Using MACS2 peaks called from H3K27ac ChIP-seq marks as true labels, we explored dilations 1, 4, and 8 and kernel sizes 31, 41, 51, and 61 in the MCF-7 cell line data, evaluating precision, recall, and F1 across 22, excluding sex, chromosomes. We observed that kernel size has little effect on precision and recall values, while dilation is influential. In particular, larger dilation improves the precision but lowers the recall, leaving F1 scores roughly unchanged (Supplementary Figure S8a).

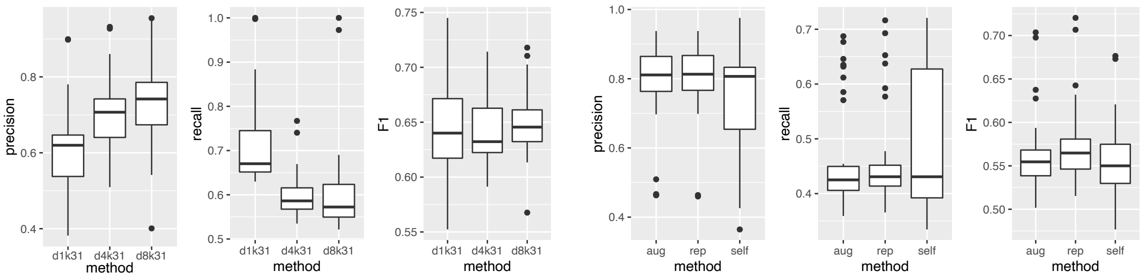


(a) (b)

Figure S8: Varying RCL hyperparameters and inputs. (a) Change in precision, recall and F1 score across 22 chromosomes at different choices of dilation in $\{1, 4, 8\}$ for the MCF-7 cell line data. d$n$ means dilation is $n$, k31 means kernel size is fixed at 31. (b) Change in precision, recall and F1 score for different choices of the contrasted data across chromosomes. Abbreviations: "aug", method that contrasts the coverage vector with the reversed coverage vector, "rep", method that contrasts the coverage vector with the coverage vector of a second replicate, "self", method that contrasts the coverage vector with itself.

##### S3.1.2 Ablation study: contrasting biological replicates improves method performance

To study the value of replicates, we contrasted the coverage vector from the first replicate in the MCF-7 data with entities other than the coverage vector of the second replicate. Specifically, in equation (1), $\boldsymbol{x}_{r'i}$ is replaced with $\boldsymbol{x}_{ri}$ (self) or $\boldsymbol{x}_{ri}$ in reversed order (augmentation). Contrasting to the coverage vector of the second replicate gives the best performance overall, but contrasting with the augmentation formed by reversing the first coverage vector has only slightly worse performance (Supplementary Figure S8b). Thus, a valid approach for using RCL when no biological replicates are available might be to create viable augmentations of the single coverage vector. On the other hand, contrasting the coverage vector of the first replicate to itself resulted in the worst performance and notably increased the variation in performance across the different chromosomes. Therefore, contrasting coverage vectors from different biological replicates during learning helps the model retain essential peak information and exclude distracting noise information to improve overall performance.

###### S3.1.3 Training vs. testing performance: little evidence of overfitting

RCL is an unsupervised method, so we fit the model to all the provided data, or a random subset if the data is large, then predict on all the provided data. While we are concerned that developing the model on MCF-7 data may limit the generalizability of RCL to other datasets, we do not need to separate training and testing data within a given dataset to guarantee no model overfitting. Nevertheless, RCL is sufficiently complex that it may overfit a particular set of segments. We can use training and testing datasets to assess whether the RCL model is overfit if it has decreased performance on testing data. We randomly sampled 80% as the training data and predicted on the remaining "test data" in 10 replicated experiments. The performance is assessed across all chromosomes. The average precision and recall for the training set are 0.741 and 0.593, while the average precision and recall for the testing set are 0.731 and 0.602, representing a change of less than 2% (average absolute change for precision and recall are 0.01 and 0.02). There is no evidence of overfitting in the MCF-7 data. Further testing on four independent datasets confirms that the RCL advantage over other methods is robust across datasets, despite the model development done on MCF-7.

##### S3.2 Score distribution analysis shows clear separation between peak and non-peak regions predicted by RCL

To understand the connection between peak predictions and the scores assigned by each method, we analyzed the score distribution of peaks called within the ChromHMM labeled regions. In particular, we examined scores from MACS (`-q 0.05`), ChIP-R (`-a 0.05`), LanceOtron, HMMRATAC and RCL-MED, for MCF-7, A549, and K652, or RCL-C2 for the GM12878 and mouse placenta data using pairs plots (Figure S9). In general, we observed that MACS and ChIP-R scores are highly correlated, likely because MACS and ChIP-R scores were computed from the same input regions called on individual replicates by MACS (see Methods). RCL scores tend to be least correlated with scores of the other methods, suggesting RCL is detecting distinct signals of peaks. Overall, the separation between peak and non-peak scores is more obvious for RCL than other methods, though the deep learner LanceOtron also shows strong separation. For example, some predictions are exceptionally confident for RCL-C2 (absolute logit-transformed scores $> 10$) and LanceOtron (absolute logit-transformed scores $> 4$) on the MCF-7 data (Figure S9a). Interestingly, RCL also has many intermediate scores (logit transformed scores near 0), where the separation between peaks and non-peaks is less obvious. We calculated PRAUCs using just these intermediate-scoring regions for MCF-7, and RCL-C2 still achieves better separation (PRAUCs: 0.718, 0.483, 0.492 and 0.480 for RCL-C2, MACS, HMMRATAC and CHIP-R, respectively), indicating it makes better predictions even in challenging regions.

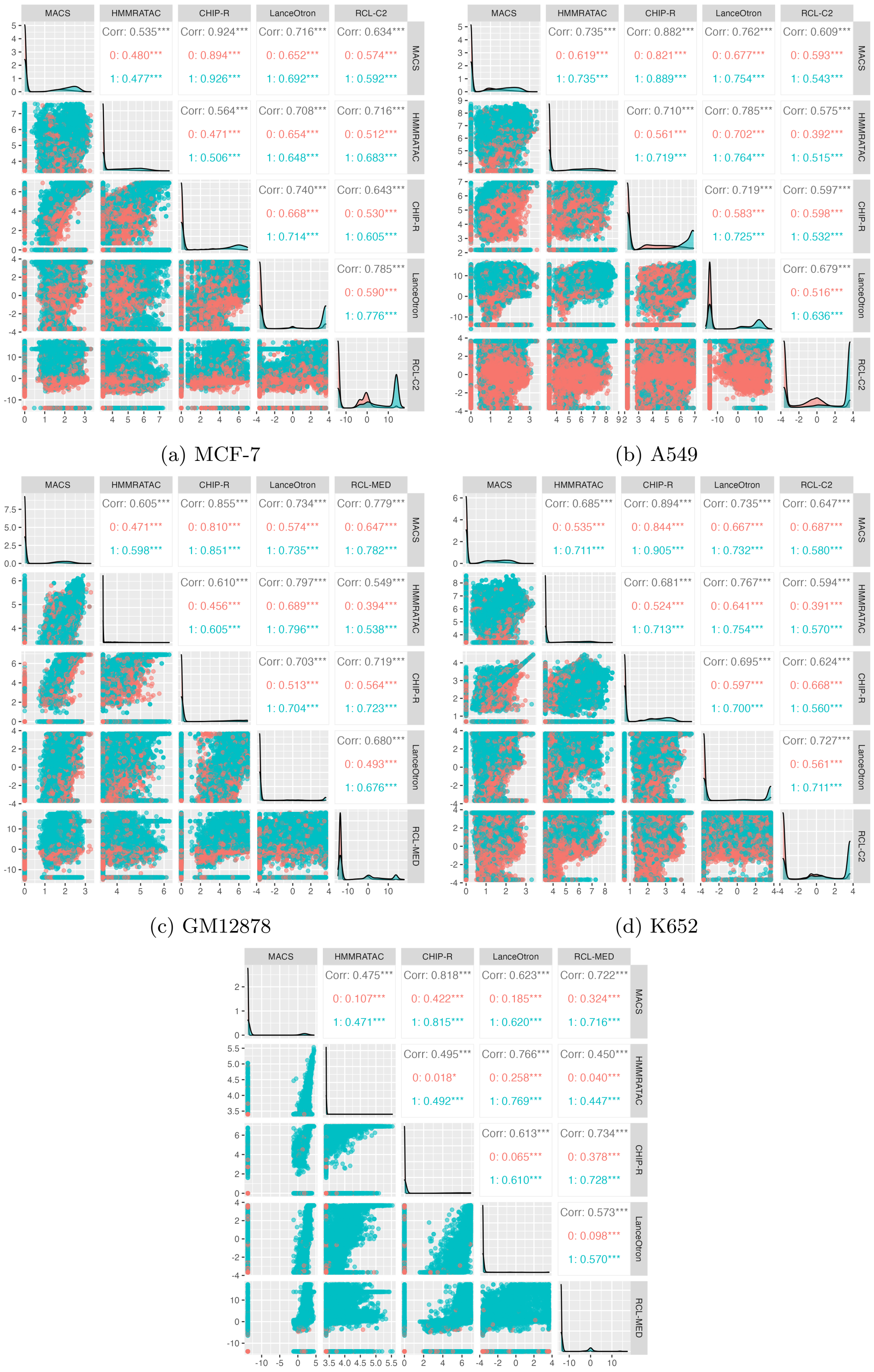


(a) MCF-7

(b) A549

(c) GM12878

(d) K652

(e) Mouse placenta

Figure S9: Pairs plots of MCF-7 (a), A549 (b), GM12878 (c), K652 (d) and mouse placenta (e) data. Pink (0, upper triangle of each pairs plot) indicates non-peaks as determined by Chrom-HMM and blue (1) indicates peak. RCL-C2 and LanceOtron scores are logit transformed, others are log transformed.

#### S3.3 RCL predictions are stable with more replicates and high coverage replicates

To assess how the number of replicates affects RCL performance, we supplied increasing numbers of biological replicates to RCL in the A549 (three total available replicates) and GM12878 (four replicates) data and used ChromHMM labels to evaluate the performance. Of note, one GM12878 replicate has higher library size than the others (64.8 million reads compared to an average 20.2 million reads in the other three replicates). When adding GM12878 replicates, we added the high library replicate last. RCL performance hardly changed with additional replicates, even the replicate with high coverage (Figure S10). It is possible that in these data, replicates are so similar that additional replicates provide no new information to the contrastive learner. Since the datasets we investigated were of high quality, it remains to be seen if particularly noisy replicates can affect RCL performance. Based on our experiments, however, we recommend using all available replicates. It does not hurt, hardly takes much more computational time (see Discussion), and could help RCL call peaks.

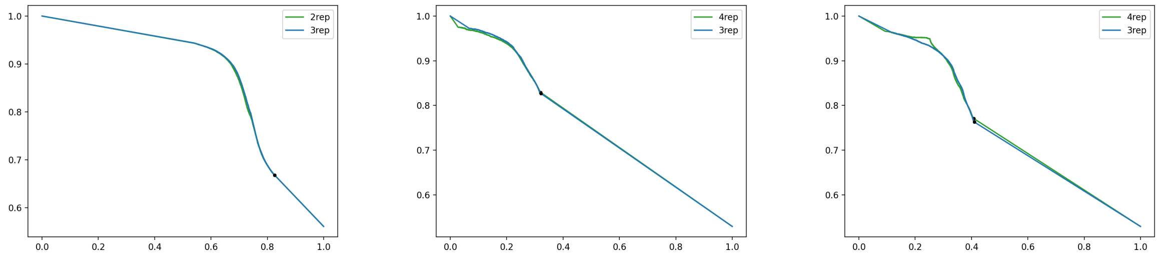


(a) A549. (b) GM12878 ($-t$ 2). (c) GM12878 ($-t$ median).

Figure S10: RCL performance when supplying additional replicates, using the A549 (a) and GM12878 (b), (c) datasets. $n$rep means RCL is provided with $n$ replicates.

#### S3.4 Extent of candidate peak regions $\mathcal{A}$ are crucial to RCL success

Peak calling involves identifying likely peaks and their boundaries. Methods are indirectly judged on their ability to identify region boundaries since evaluations typically check if the predicted peaks overlap with labeled regions (see Methods, (Tarbell and Liu 2019), (Hentges et al. 2021)). To analyze the role of the candidate regions in $\mathcal{A}$ we pass to RCL for training and prediction (see Results §1.1.1), we conducted several experiments on the MCF-7 data (here, using $t = 2$, RCL-C2) and A549 (here, using default $t$, RCL-MED). The first set of experiments explored the role of the RCL score in its performance on MCF-7. We tried scoring the RCL candidate regions with the mean coverage across replicates (denoted as PRE in Figure S11a), and we mapped other methods' predictions to the RCL candidate peak regions in $\mathcal{A}$, as we had for RCL segment scores: if multiple peaks overlap one candidate region, the region was scored with the weighted mean score of the overlapping peaks (see Methods). For ChIP-R and MACS, we used the predictions and scores from runs with $q$-value $= 0.5$. The second set of experiments explored the role of the candidate regions $\mathcal{A}$ in RCL performance on A549. Instead, we used the 1,000 bp segments as the prediction regions. We also attempted to use the decoder output $\hat{\boldsymbol{m}}_{ri}$ to identify peak centers. Specifically, we identified all sites with denoised coverage above the 0.5 quantile within the 1,000 bp segment, merged adjacent sites, and used the largest merged region as the prediction region.

Candidate peak regions in $\mathcal{A}$ scored simply by average coverage performed better than all other competing methods except RCL-C2, where scores came from the contrastive learner (Figure S11a). These experiments confirm that high coverage is highly predictive of true peaks, and our method to choose candidate peak regions is effective. However, the experiments also showed that other signals extracted by the contrastive learner contributed substantively to the best peak predictions. However, using these superior RCL scores on other prediction regions, either the 1,000 bp segments themselves or an approximation of the peak center estimated from the decoded signal, performed abysmally (Figure S11b). Further investigation (data not shown) indicated that signal from strong, narrow peaks in 1,000 bp regions leaked false positive calls into nearby non-peak regions. However, our attempt to shrink down 1,000 bp segments to peak centers performed even less well. The difficulties revealed by these experiments stem from the fact that RCL is not determining the peak extent. Further discussion on how an unsupervised learning method could accomplish both facets of peak calling, presence and extent, are mentioned in the main text Discussion.

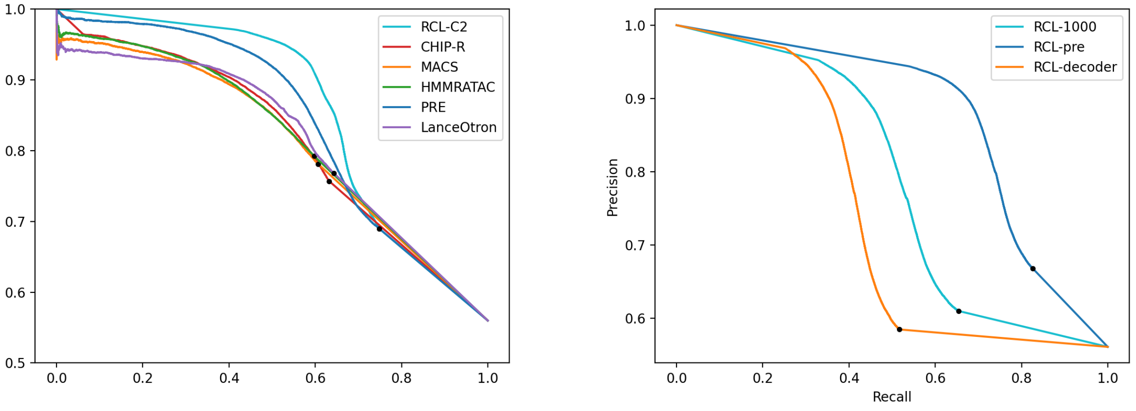


(a) MCF-7 varying score

(b) A549, varying region

Figure S11: PR curve when varying prediction region or prediction score. (a) We vary the score assigned to MCF-7 candidate regions in $\mathcal{A}$. PRE represents the candidate peak regions scored with mean coverage across replicates; ChIP-R, MACS, HMMRATAC, and LanceOtron are the respective method predictions mapped candidate peak regions in $\mathcal{A}$; RCL-C2 are the peak predictions using the full RCL framework, duplicate of the same curve in Results Figure 2a. (b) We substitute prediction regions other than the candidate regions in $\mathcal{A}$ in A549. RCL-1000 uses 1,000 bp segments as prediction regions. RCL-pre uses the candidate regions defined in $\mathcal{A}$, duplicate of the same curve in Results Figure 2d. RCL-decoder uses subregions within the 1,000 bp segments obtained from the decoded coverage (see text).

#### S3.5 Non-default settings of MACS may result in a better performance

By default, MACS estimates a shift and window size from paired end read data to shift the read alignment positions before counting the number of reads aligning in windows of the given size. This procedure is motivated by the bimodal patterns of read coverage seen in transcription factor (TF) ChIP-seq data. This same procedure is often applied to call peaks on ATAC-seq data, but shifting may be unnecessary when open chromatin regions are not occupied by any TF. Researchers specifically interested in the Tn5 cutting sites can impose a biologically defined shift and window size instead of using the estimation from data (see `https://github.com/macs3-project/MACS/blob/master/docs/callpeak.md`), but generally there is no consensus on the best way to use MACS to call ATAC-seq peaks. As a result, we used default options throughout this manuscript. However, we also explored a non-traditional setting, where the shift size is 0 base pair (bp) (`--shift 0`), window size is 2 bp (`--ext-size 1`), and $q$-value is set at a relaxed threshold (0.5), hereafter the "shift0" settings.

Interestingly, MACS "shift0" led to improved performance of the peak prediction for high confidence (low recall, high precision) peaks, even better than RCL on the MCF-7 dataset for this region of the PR curve (Figure S12). This setting calls a *large* number of peaks. For example, MACS "shift0" predicted 10,963,599 peaks, over 60% of which were 1 bp long, for the A549 data. MACS "shift0" may easily predict single sites in high confident peak regions, where coverage is clearly higher than the background. However, the model suffers from high false positive rates when trying to predict more difficult peaks. Despite the excellent performance in strong peak regions, the very narrow peaks may be difficult to use in downstream analyses or interpret biologically. Furthermore, it is clear that peak calling for ATAC-seq data with ChIP-seq tools remains imperfect and unpolished.

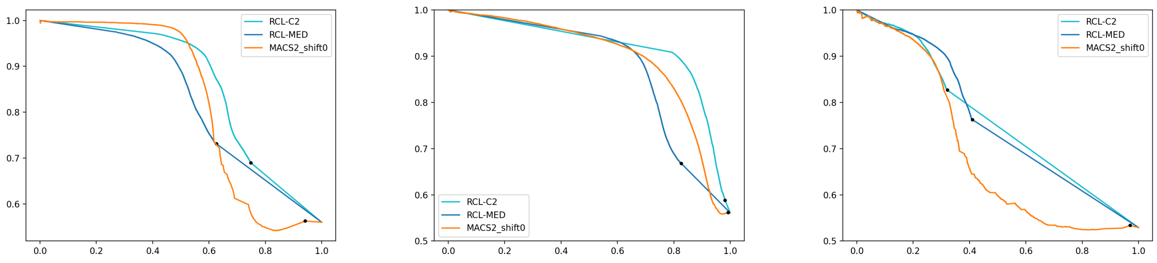


(a) MCF7 (b) A549 (c) GM12878

Figure S12: PR curves when comparing non-default MACS settings' results with RCL. MACS2_shift0 represents no shifting reads and the window is 2bp.
